# Corpus callosum morphology across the lifespan in baboons (*Papio anubis*): a cross-sectional study of relative mid-sagittal surface area and thickness

**DOI:** 10.1101/2020.12.07.414367

**Authors:** René Westerhausen, Adrien Meguerditchian

## Abstract

The axons forming the corpus callosum enable integration and coordination of cognitive processing between the cerebral hemispheres. In the aging human brain, these functions are affected by progressive axon and myelin deteriorations, which results in a substantial atrophy of the midsagittal corpus callosum in old age. In non-human primates, these degenerative processes are less pronounced as previous morphometric studies on capuchin monkey, rhesus monkeys, and chimpanzees do not find old-age callosal atrophy. The objective of the present study was to extend these previous findings by studying the aging trajectory of the corpus callosum of the olive baboon (*Papio anubis*) across the lifespan. For this purpose, total relative (to forebrain volume) midsagittal area, subsectional area, and regional thickness of the corpus callosum was assessed in 91 male and female animals using non-invasive MRI-based morphometry. The studied age range was 2.5 to 26.6 years, and the sample included 11 old-age animals (above the age of 20 years). Fitting lifespan trajectories using general additive modelling (GAM) we found that the relative area of the total corpus callosum and the anterior subsection follow a positive linear trajectory. That is, both measures increased slowly but continuously from childhood into old age, and no stagnation of growth or decline was observed in old age. Thus, comparable with all other non-human primates studied to-date, baboons do not show callosal atrophy in old age. This observation lends supports to the notion that atrophy of the corpus callosum is a unique characteristic of human brain aging.

---

The corpus callosum as the major white-matter commissure, is a critical channel for the integration of information (e.g., Steinmann et al., 2018; Westerhausen, Gruner, Specht, & Hugdahl, 2009) and coordination of processing in the two cerebral hemisphere (e.g., Davis & Cabeza, 2015; Thiel et al., 2006). Thus, it appears little surprising that human neuroimaging studies report an association of individual differences in corpus callosum morphology with differences in higher cognitive abilities (e.g., Danielsen et al., 2020; Dunst, Benedek, Koschutnig, Jauk, & Neubauer, 2014; Hulshoff-Pol et al., 2006; Luders et al., 2007; Westerhausen et al., 2018). In the aging brain, however, these functions of the corpus callosum are affected by a progressive degeneration of callosal axons, evidenced by a reduction in number and density of small myelinated axons (Fan et al., 2019; Hou & Pakkenberg, 2012; Køster, Jesper, & Bente, 2018; Lynn et al., 2020). Diffusion MRI studies, likely reflecting these axonal alterations, find a decrease in fractional anisotropy and an increase in radial diffusivity in older age (e.g., Hasan et al., 2009; Ota et al., 2006; Pietrasik, Cribben, Olsen, Huang, & Malykhin, 2020; Skumlien, Sederevicius, Fjell, Walhovd, & Westerhausen, 2018). Morphometric analyses of the midsagittal corpus callosum additionally report a substantial atrophy of callosal surface area (Doraiswamy et al., 1991; Hasan, Ewing-Cobbs, Kramer, Fletcher, & Narayana, 2008; Prendergast et al., 2015; Salat, Ward, Kaye, & Janowsky, 1997; Skumlien et al., 2018) and thickness measures (Danielsen et al., 2020). This atrophy of the corpus callosum is stronger than what would be predicted from the parallel ongoing decline in overall forebrain and white-matter volume, suggesting a progressive decline in structural callosal connectivity in the aging brain (Danielsen et al., 2020; Salat et al., 1997; Skumlien et al., 2018). Furthermore, such a decline was found accentuated in patients with Alzheimer’s dementia in comparison to controls (e.g., Frederiksen et al., 2011; Thomann, Wüstenberg, Pantel, Essig, & Schröder, 2006; Wiltshire, Foster, Kaye, Small, & Camicioli, 2005), underlining the potential contribution of callosal decline to changes in cognitive abilities in aging humans.

To reach a better understanding of the aging corpus callosum in humans, it is relevant to determine whether this over□ proportional callosal decline is a specific human trait, or represents a common feature of brain aging across species. To answer this question, comparative approaches with non-human primates promise to be most informative for identifying human□ specific brain aging characteristics (Colman, 2018; Rilling, 2014). To date, while only for a few primate species morphometric studies of the aging corpus callosum exist, the available findings seem to be consistent. In both capuchin monkey (*Cebus paella*; Phillips & Sherwood, 2012) and rhesus monkeys (*Macaca mulatta*; Bowley, Cabral, Rosene, & Peters, 2010; Lacreuse et al., 2005) no significant old-age reduction of midsagittal area was found. Likewise, in chimpanzees (*Pan troglodytes*) a plateau or continuous increase of relative corpus callosum area and thickness has been reported in old age (Hopkins et al., 2016; Hopkins & Phillips, 2010; Westerhausen et al., 2020). A recent direct comparison of human and chimpanzee lifespan trajectories, additionally confirms that the old-age reduction in total midsagittal callosal area is significantly stronger in humans (Westerhausen et al., 2020). Thus, taken together, old-age atrophy of the corpus callosum has not been reported in any other primate species but humans, so that it appears tempting to conclude that it represents human□ specific phenomenon of brain aging. However, the analyses of further primate species is warranted to substantiate this conclusion, and the comparison of the aging trajectories across different species might offer insight into the reasons for interspecies differences.

Following the above, the primary aim of the present study was to determine the aging trajectory of corpus callosum development in the olive baboon (*Papio anubis*) and, by doing so, extending the available literature. To this end, we analysed the corpus callosum of 91 animals using non-invasive MRI□ derived measures of total relative (to forebrain volume) midsagittal area, subregional area, and regional thickness. The sample covered an age range from 2.5 to 26.6 years, and included 11 old-age animals above the age of 20 years (approximately equivalent to a human age of 60-65 years, see e.g. Franke et al., 2017; Havill et al., 2005). Age trajectories were fitted applying general additive modelling (GAM, Wood, 2017). As the sample includes animals from childhood to old adulthood, we expected an increase in relative callosal area and thickness into adulthood but – following the available literature summarized above – no decline in old adulthood. Thus, a positive linear or saturation curve was predicted to describe the relationship of age and the callosal measures. Additionally, we examined a possible sexual dimorphism in the corpus callosum, and explored potential differences in the age trajectories between the sexes.

## Material and Methods

### Sample

The structural MRI of 96 olive baboons (*Papio anubis*) were retrieved from the database of the Laboratoire de Psychologie Cognitive (UMR7290, Marseille, France). The *in vivo* MRI were initially collected between August 2013 and January 2015 (e.g., Marie et al., 2018) from baboons housed in south of France at the Station de Primatologie CNRS (UPS846, Rousset). Of the total dataset, for two juvenile females, the data of birth was not available, so that the exact age at time of scanning could not been determined. Three additional datasets (1 juvenile, 1 adult male, 1 old female) were too noisy to achieve a good segmentation quality of the corpus callosum (see below). The remaining 91 animals (57 females, 34 males) had a mean age of 11.7 ± 5.9 years and covered an age range from 2.5 to 26.6 years.

### MRI acquisition

Magnetic resonance imaging (MRI) was conducted on 3T scanner (MEDSPEC 30/80 ADVANCE, Bruker) located at the Marseille MRI Center (Institut de Neuroscience de la Timone). T1-weighted brain images were acquired using either of two MPRAGE sequences. The imaging parameters of the two sequences were identical (TR: 9.4 ms; TE: 4.3 ms; flip angle: 30°; inversion time: 800 ms), while the field of view (FOV) of the sequences differed to accommodate differences in head size between animals (sequence 1: 108 × 108 × 108 mm; isotropic voxel size: 0.6 mm^3^; sequence 2: 126 × 126 × 126 mm, and isotropic voxel size: 0.7 mm^3^).

The MRI procedure has been described elsewhere in detail (Love et al., 2016). In brief, animals were premedicated at the Station de Primatologie by intramuscular injections of ketamine to facilitated transport to the MRI facility. Anaesthesia was maintained during scanning using drip irrigation setup including tiletamine, zolazepam and NaCl, while the animals’ cardiovascular and respiratory parameters were under constant monitoring. After conclusion of the imaging session, the animals remained under surveillance to assure they were fully awake and had recovered from anesthesia in good health, before the animals were returned back to their social group.

The experimental procedure complied with French laws and the European directive 86/609/CEE and has been approved by the ethic committee of Provence (Agreement number for conducting experiments on vertebrate animals at the Station de Primatologie CNRS: D130877).

### Corpus callosum segmentation and raw measurements

Midsagittal surface area and thickness measures were determined based on white-matter segmentations of T1-weighted images in native space as obtained using SPM12 routines (Statistical Parametric Mapping, Wellcome Department of Cognitive Neurology, London, UK) and utilizing the Haiko89 template (Love et al., 2016). The segmentation was followed by a rigid-body co-registration (i.e., preserving size and shape of the corpus callosum) of the resulting images to the respective T1-template to achieve a non-tilted midsagittal plane.

The structure of the corpus callosum was then automatically identified on the individual midsagittal white-matter maps of each individual and visually inspected, using the same routines previously described for human data (Westerhausen et al., 2016). That is, manual correction were applied where necessary using in house graphical user interface programmed in Matlab (MathWorks Inc. Natick, MA, USA). For example, when voxels belonging to the fornix were fused with the corpus callosum body, they were manually removed. In a next step, the tip of the rostrum (defined as the posterior-most voxel of the in-bend anterior callosal half) and the base of the splenium (ventral-most voxel in the posterior half) were identified. Then, the callosal mask was rotated to achieve horizontal orientation of the imagined line connecting rostrum tip and base of the splenium. The rotated mask of each individual was the basis for both area and thickness measures. The quality of each individual segmentation was then rated on a three□ point scale (0 = not usable, 1 = low, but acceptable quality, 2 = good quality) and only segmentations rated 1 and above are included in the present analyses.

The size of the callosal mask multiplied with the voxel dimensions in the sagittal plane represented the total midsagittal area in mm^2^. To determine subregional area measures, the mask was divided in thirds relative to anterior-posterior length of the structure (see Supplementary Figure S1). Related to the frequently applied subdivision schema suggested by Witelson (Witelson, 1989) for humans, the anterior third covers genu and rostrum of the corpus callosum, the middle third includes the truncus regions, and a posterior third contains isthmus and splenium.

For measuring callosal thickness, an outline of the midsagittal surface was automatically generated from the individual callosal masks. Rostrum tip and base of the splenium points were then used to divide the total outline into a ventral and dorsal part. A midline between ventral and dorsal outline was created as geometric average of 100 corresponding support points spaced equidistantly on the two outlines. In a next step, 60 equidistant measurement points were placed on this midline (see Supplementary Figure S1). Callosal thickness was then determined as the distance between ventral and dorsal outline measured orthogonal to the midline at these measurement points. The number of 60 points was chosen as it guarantees a good sampling of the structure of corpus callosum while not inflating the number of statistical tests excessively (Westerhausen et al., 2016). It can also be seen as compromise between previously used 29 to 100 points (Clarke, Kraftsik, Van der Loos, & Innocenti, 1989; Luders, Narr, Zaidel, Thompson, & Toga, 2006).

Finally, in order to account for differences in brain size, all area and thickness measures were divided by forebrain volume (FBV, see next paragraph) following the approach suggested by Smith (Smith, 2005). That is, area measures were divided by FBV raised to the power of 2/3 (i.e., FBV^0.666^) and thickness measures were divided by FBV to the power of 1/3 (i.e., FBV^0.333^). This approach was chosen, as only if FBV is expressed in the same unit as the respective callosal measure, the resulting ratio is expected to be stable if corpus callosum and FBV change proportional to each other (Smith, 2005). Any positive or negative deviation from a stable ratio consequently indicates an over-proportional growth and decline of the corpus callosum, respectively. The resulting ratios are henceforth referred to as relative area and relative thickness measures.

### Forebrain-volume estimation

FBV was determined using an inverted-mask approach. Firstly, gray□ and white□ matter maps were created for each individual in native space using SPM segmentation routines and the Haiko89 baboon template (Love et al., 2016). Then, a custom forebrain mask covering the supratentorial brain was transferred from Haiko89 standard space to the individual brain by using the same transformation-inversion parameters used when creating the tissue segmentations in native space. Finally, individual FBV was estimated as the sum of gray□ and white□ matter probabilities within the mask in native space. Thus, the resulting FBV estimates did not include CSF compartments. FBV was preferred over estimates of total intracranial volume, as it mostly includes the volume of the cerebral hemispheres, which are connected via the corpus callosum, while it excludes brain structures (e.g., brain stem, cerebellum) which are not known to have callosal axons (Schmahmann & Pandya, 2006).

Of note, the correlation of FBV with total corpus callosum area was r = 0.40 (t = 4.07, df = 89, p < 0.001) underlining the necessity of considering differences in FBV when analyzing the aging trajectories of the corpus callosum.

### Statitsical analysis

The age trajectories of relative total and subregional area as well as of relative thickness measures were fitted using generalized additive modelling (GAM) with the “mgcv” R-package (v1.8□ 31; (Wood, 2017); R build: 3.6.2). A cubic regression splines with 4 knots as a basis dimension was evaluated to be appropriate for all analyses. The animal’s Sex was included as covariate in all analyses to test for sexual dimorphisms (irrespective of age). In a second step, follow-up analyses were conducted to test for sex differences in the aging trajectories. That is, the GAM analysis was set up to model the trajectory for females, and test for a deviation of the male from the female trajectory. In all analyses, the models were fitted using restricted maximum likelihood (REML) estimates. Example R code is provided in Supplement Section 2. Effect size measures, expressed as proportion of explained variance (ω^2^), were provided for all analyses. For the segment□ wise analyses of relative thickness, p□ values were adjusted to yield a false discovery rate (FDR) of 5%.

### Data availability statement

The tabulated data and analysis scripts supporting the present analyses are available on the OSF platform (https://osf.io/5yzb3/). The raw MRI data is available on request from Adrien Meguerditchian.

## Results

### Midsagittal surface area

The mean outline across individual is presented in Supplementary Figure S1. The mean total corpus callosum area of the sample was 96.9 ± 15.4 mm^2^ (range: 57.9 to 143.8 mm^2^), and mean relative area was 0.039 ± 0.006 (range: 0.023 to 0.059).

For the total corpus callosum, a significant linear effect (edf = 1.00) of Age was found on relative area (*F* = 5.32, *p* = .023, ω^2^ = .045; see Fig. 1). Further analyses by corpus callosum subsection showed that the positive linear association (edf = 1.00) with age was significant only in the anterior third section (*F* = 5.64, *p* = .020) which explained 4.9 % of variance in the data. For middle third (edf = 1.00, *F* = 3.54, *p* = .063, ω^2^ = .027) and posterior third sections (edf = 1.56, *F* = 1.89, *p* = .11, ω^2^ = .020) the age effect was not significant.

**Figure 1.**
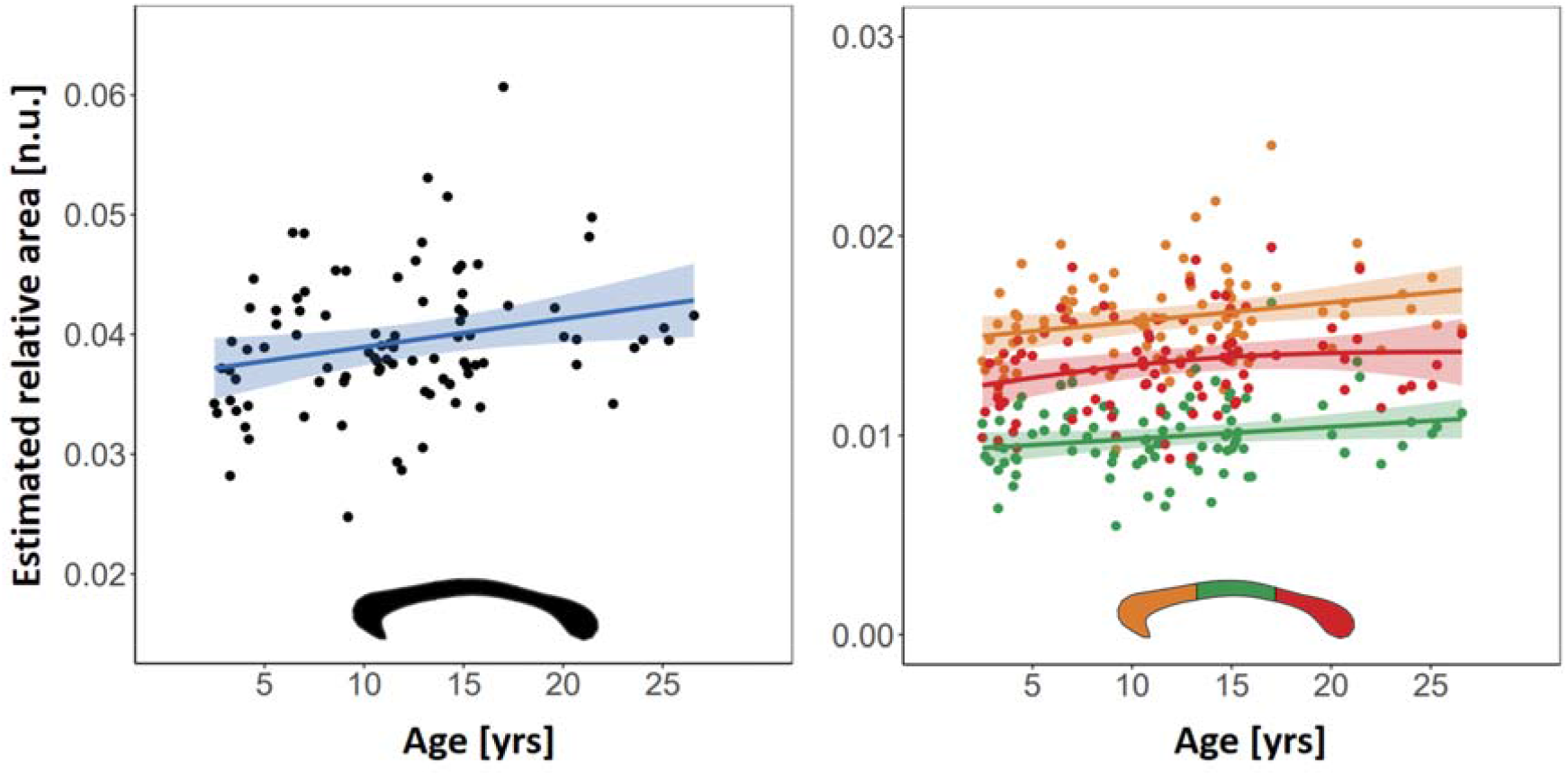
Lifespan trajectories of relative corpus callosum area. The graphs show the GAM fitted age trajectories (shaded area 95% confidence bands) corrected for Sex. Left panel: total relative area, right panel: subsection relative area (colour-coded by subsection, as indicated by inlay). Of note, the ratio (i.e., relative area) does not have a unit (n.u.) as the units of numerator and denominator cancel out.

No significant main effect of sex was found for neither the total relative corpus callosum area (*t*(88.0) = -1.13, *p* = 0.26, *ω*^2^ < .01) or any of the three subsections (anterior third: *t*(88.0) = -0.51, *p* = 0.61, *ω*^2^ < .01; middle third: *t*(88.0) = -1.78, p = 0.08, *ω*^2^ < .01; posterior third: *t*(87.4) = -0.83, p = 0.41, *ω*^2^ < .01). The follow-up analysis testing for sex differences in the age trajectories did not reveal any significant differences. That is, the deviation of the male trajectory from the female trajectory was neither significant for total relative callosal area nor for any of the three subregions (all F <1.46, all p > .27, all *ω*^2^ < .02).

For matter of completeness, we repeated the above analyses using absolute instead of relative area as dependent variable. As can be seen in Supplement Section 3, these analyses mirrored the findings observed when analysing the relative measures.

### Regional thickness

The segment-wise thickness analysis did not reveal any significant age trajectory fits after FDR adjustment (all edf < 1.98, all *p*_FDR_ >.83; all *ω*^2^ < .11; see supplement Table S1 for test statistics by segment). Exploring uncorrected p-values, 6 of the 60 segments showed p-values below an alpha of 5%, of which 5 were in close proximity of each other in the posterior corpus callosum (*ω*^2^ between .06 and .11; see Fig. 2). The aging trajectories in this cluster were characterised by a saturation function (edf between 1.68 and 1.92): a continuous increase from childhood to mid adulthood which is followed by stable thickness into old age.

**Figure 2.**
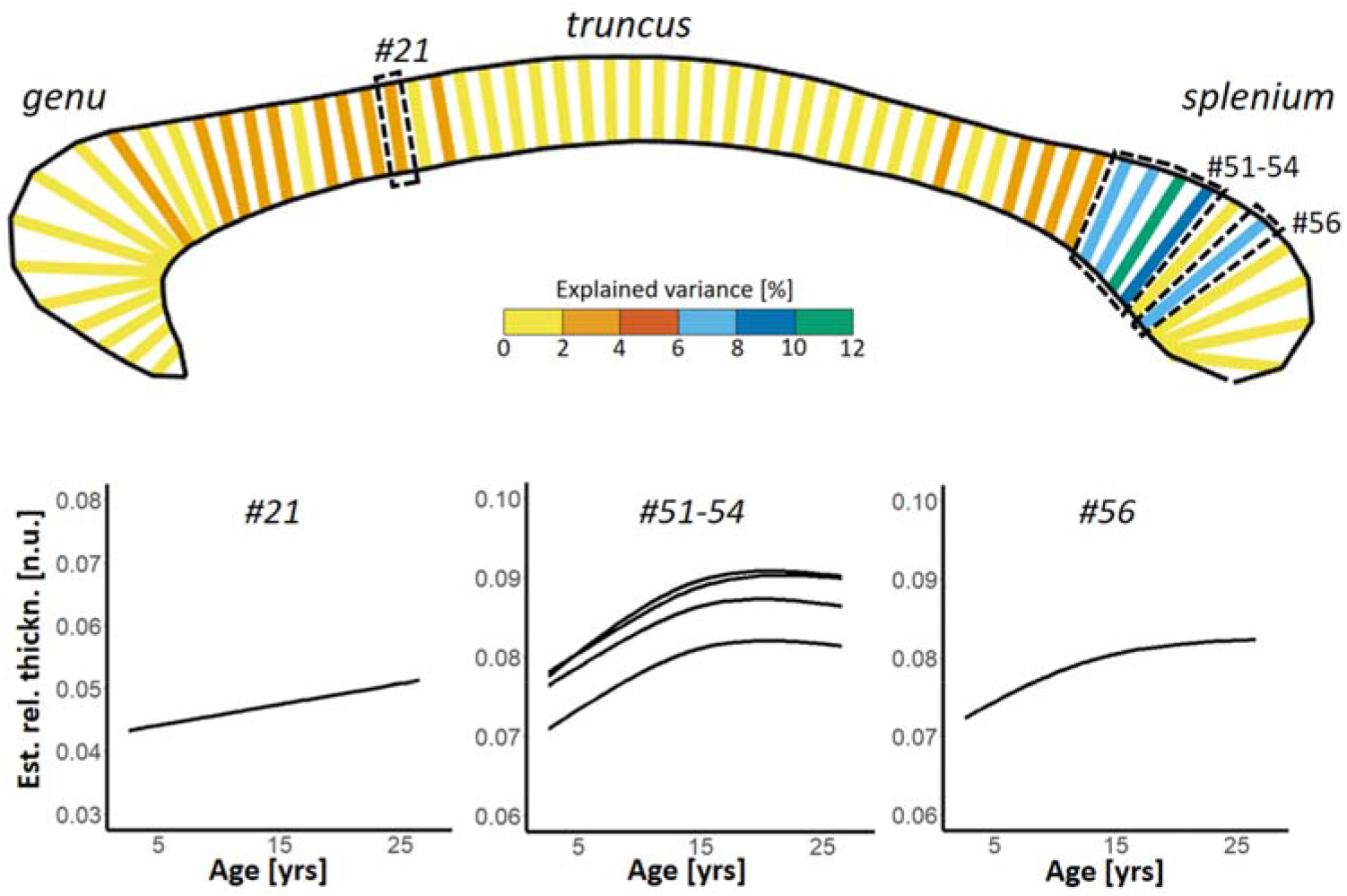
Results of the segment-wise analysis of the Age effect on relative thickness. The upper panel color-codes the proportion explained variance of the Age effect for each segment within the mean outline of the baboon corpus callosum. For neither of the segments the Age effect was significant after FDR correction. The highlighted segments (dotted lines) show as significant effect when considering uncorrected p-values (# segment number). The graphs in the lower panel show the GAM fitted lifespan trajectories for these segments corrected for Sex. Of note, relative thickness does not have a unit (n.u.) as the units of numerator and denominator cancel out.

Likewise, no main effect auf Sex was found at an FDR of 5% (all *p*_FDR_ >.72; all *ω*^2^ < .08), nor did the analysis yield any significant differences in the age trajectories between sexes (all *p*_FDR_ >.07; all *ω*^2^ < .20).

Repeating these analyse for absolute (rather than relative) thickness measures also did not yield any significant Age or Sex effects after FDR correction (see Supplement Section 3 for details).

## Discussion

In the present study, we examined age- and sex-related difference in the morphology of the baboon corpus callosum. The main finding was that relative area of total corpus callosum follows a linear lifespan trajectory. That is, midsagittal area increases slowly but continuously from childhood into old age. In this, the present study supplements the findings of a series of previous studies suggesting that the aging trajectories of the corpus callosum differ substantially between primate species. The here reported linear trajectory of baboons, best resembles the rather flat lifespan trajectory found in capuchin monkeys (Phillips & Sherwood, 2012), but appears dissimilar to the more accentuated trajectories found in chimpanzees (Hopkins et al., 2016; Hopkins & Phillips, 2010; Westerhausen et al., 2020) and humans (Danielsen et al., 2020; Rauch & Jinkins, 1994; Salat et al., 1997). Figure 3 was created to follow-up on this comparative perspective. It provides a visual comparison of GAM-fitted lifespan trajectory of the total baboon corpus callosum with the trajectories found in humans and chimpanzees reported in a previous study (Westerhausen et al., 2020). Adjusting for differences in longevity between the three species (Figure 3, left graph) it gets obvious that the trajectories deviate from each other both during development and in old age. That is, the slope of the trajectory of baboons is less steep than the trajectories of the two other species during childhood and young adulthood. In old age, while the corpus callosum appears to increase continuously with age in baboons, chimpanzees show a plateau, and humans show a rapid decline in corpus callosum area.

**Figure 3.**
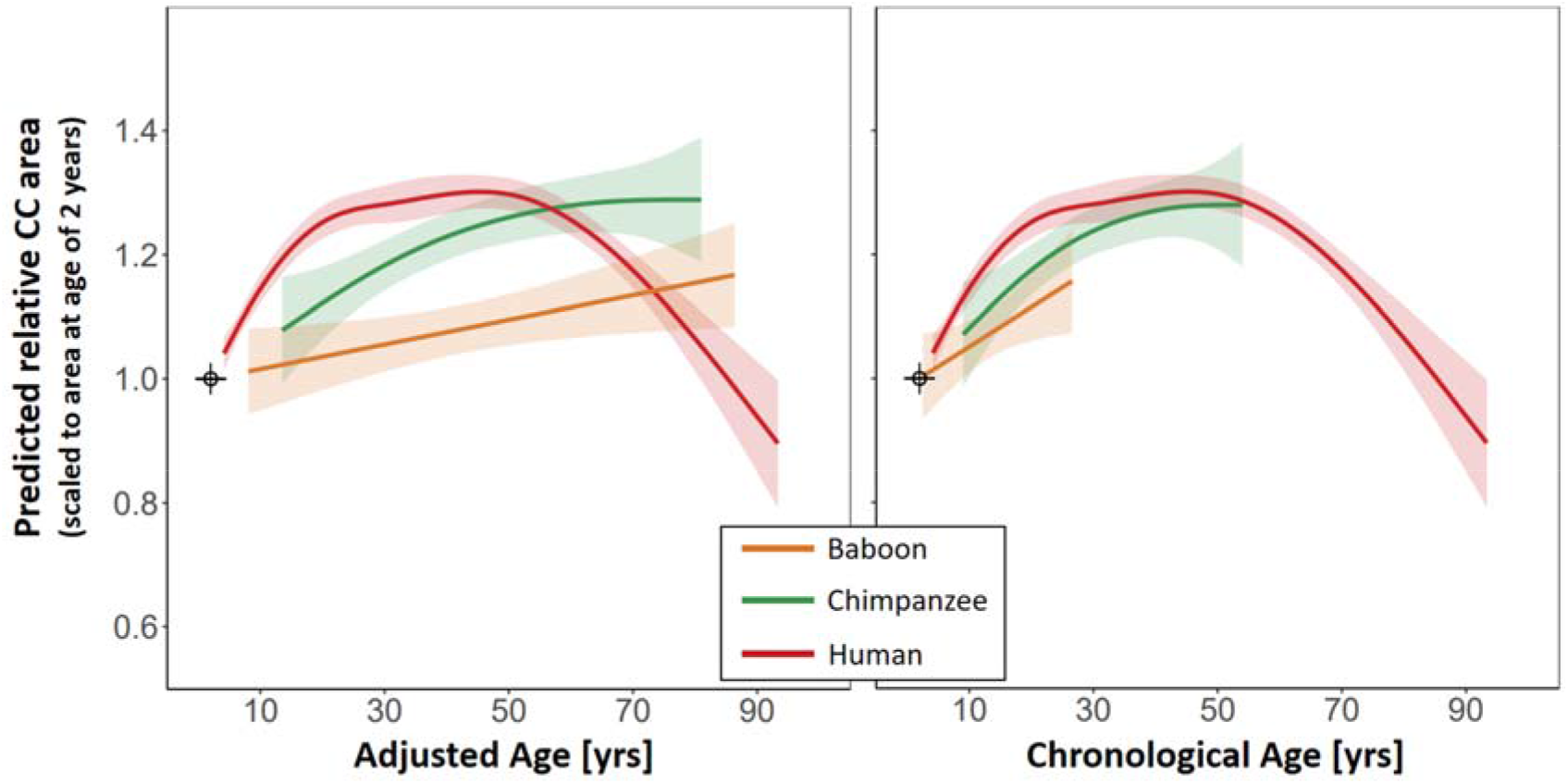
Comparison of lifespan trajectories of the corpus callosum between baboons, chimpanzees, and humans. The graphs shows the fitted GAM trajectories for each species (shaded area represent 95% confidence bands). The left panel compares the trajectories after adjusting for differences in longevity between the species; i.e. the age of the baboon and chimpanzees were transferred to a human age equivalent (referred to as “adjusted age”). The right panel shows the same comparison without age adjustment (i.e., chronological age). For both graphs, the data of each species was divided by the relative corpus callosum area predicted for the age of 2 years using the fitted GAM model of the respective species. Thus, a value of 1 represent the predicted relative area at the age of 2 years. For more details regarding the graphs please refer to Supplement Section 5.

While these species differences in the callosal lifespan trajectory appear striking, it remains matter of speculation what microstructural alterations underlie these differences, as comparative histological studies of the corpus callosum are rare (Caminiti, Ghaziri, Galuske, Hof, & Innocenti, 2009; Phillips et al., 2015) and studies comparing age□ related axonal changes between species are missing. Previous developmental studies suggest, however, that the callosal growth seen throughout childhood and adolescence is driven mainly by changes in axon diameter and/or myelination (Clarke et al., 1989) as the number of axons is extensively pruned rather than increased during postnatal development (Innocenti & Price, 2005; LaMantia & Rakic, 1990). Thus, one might speculate that the flatter slope of the baboon developmental trajectory compared with chimpanzee and human trajectories indicates comparatively mild increase in myelin or axon diameter during childhood and adolescence. As axon myelination and diameter determine the axon conduction delay (Phillips et al., 2015; Ringo, Doty, Demeter, & Simard, 1994), one might furthermore attribute these differences to differences in brain size between these species (Rilling & Insel, 1999). In the smaller baboon brain, shorter distances between the cerebral cortices of both hemispheres have to be bridged, potentially requiring a less pronounced adjustment of the axonal conduction speed during development than the larger brained chimpanzees and humans. This interpretation would also be in line with the observation that primate species with larger brains also tend to exhibit a relatively larger corpus callosum area (Manger, Hemingway, Haagensen, & Gilissen, 2010) potentially reflecting a (partial) compensation for the longer distance between the cortices with larger diameter and stronger myelinated axons (Phillips et al., 2015).

In old age, baboons do not exhibit callosal atrophy as seen in humans but rather show a continuous increase in area into old age. This observation, however, does not exclude the existence of subtle age-related axonal changes, which might not be readily reflected in morphometric measures (Chen et al., 2013). For example, while morphometric studies of the rhesus monkey do not reveal a significant reduction of midsagittal callosal area in old animals (Bowley et al., 2010; Lacreuse et al., 2005, see also Peters & Kemper, 2012), histological analyses find both degeneration of myelin sheaths (Bowley et al., 2010; Peters & Sethares, 2002) and reduction in the axon count (Bowley et al., 2010). In-vivo diffusion-imaging studies seem to mirror these findings, by revealing a significant reduction of callosal anisotropy in old aged rhesus monkeys (Makris et al., 2007; Sridharan et al., 2014). Thus, while not excluding age-related axon alterations in the baboon’s corpus callosum, the present findings at least indicate that these alterations are not sufficiently strong to cause detectable morphometric changes. In contrast, in humans the reduction of the number of myelinated axons (Highley et al., 1999; Hou & Pakkenberg, 2012; Køster et al., 2018) is apparently so dramatic, that a substantial reduction in callosal size and thickness can be found (Danielsen et al., 2020; Doraiswamy et al., 1991; Hasan et al., 2008; Prendergast et al., 2015; Salat et al., 1997; Skumlien et al., 2018). Furthermore, it has been suggested that such species differences in brain aging may partially be attributable to differences in life expectancy (Chen et al., 2013; Sherwood et al., 2011). These authors argue, that chimpanzees and rhesus monkey show only mild signs of white-matter degeneration in old age compared with humans, as they do not get sufficiently old to exhibit substantial decline. Analogously, one might argue here that baboons – due to their shorter lifespan – simply do not get old enough to reach the plateau in corpus callosum size seen in chimpanzees and humans, let alone show the atrophy observed in older humans. Figure 3 (right panel), comparing the GAM lifespan trajectories considering chronological age (rather than adjusted age), can be seen as an illustration of this hypothesis, as one can easily imagine callosal aging to follow comparable lifespan trajectories if the lifespan of baboons would be extended to chimpanzee or human age limits. However, this hypothesis implies that aging-related changes follow a universal pattern in all primate species, an assumption that is highly speculative and requires further testing.

Beyond the above, two additional observations regarding the aging trajectory deserve further discussion: differences between callosal subsections as well as divergent findings of relative area and relative thickness analyses. That is, analyses calculated separately for the three subsections find differences in the association of age and area measures between the three sections. While the age effect was significant for the anterior third, and may be considered a “trend” in the middle section, the association was not significant in the posterior section. The anterior third, including rostrum and genu, is formed from axons likely interconnecting prefrontal cortex regions (Schmahmann & Pandya, 2006) so that it appears tempting to speculate that the accentuated association found in the genu might in particular be related to the development of higher cognitive functions governed by the prefrontal lobes (e.g., Christophel, Klink, Spitzer, Roelfsema, & Haynes, 2017; Pandya & Yeterian, 1996). However, while the anterior third appears to drive the effects found for total relative area, all three sections show comparable trajectories (cf. Fig 1) and even in the non-significant posterior third the effect explains 2% of the variance in the data.

Regarding the relative thickness analysis, for neither of the segments the age effect survived correction for multiple comparison. Further exploring the data by also considering uncorrected p-values, revealed a cluster of segments in the splenium of the corpus callosum which showed an association with age of medium to large effect sizes (between 6 and 10% explained variance). The splenium is thought to connect sensory cortices or parietal brain regions (Schmahmann & Pandya, 2006), suggesting a relationship of the aging effect to the development of sensory or attentional integration between the hemispheres (e.g., Bozzali et al., 2012; Steinmann et al., 2018). Interestingly, here the aging trajectory resembled a saturation function: a rapid increase from childhood to adulthood is followed by a period of constant thickness that reaches into old age. Taking these uncorrected findings at face value, the results of relative thickness and area analysis deviate in two ways: the age effect was found in the posterior rather than the anterior third and the shape of the aging trajectory was curvilinear rather than linear. Of note, however, from a methodological perspective area measures and thickness may theoretically vary independent of each other. Thickness estimates are usually obtained at a fixed number of measurement points placed in equidistant intervals on the outline or midline of the corpus callosum (Luders et al., 2006; Walterfang et al., 2009; Westerhausen et al., 2016). Variations in the length of midline/outline between individuals will consequently affect the spacing of the measurement points but not directly affect the thickness measurements. Thus, the divergence between area and thickness measures found in the present study, might reflect that the two measures are sensitive to different features of the morphological variability of the corpus callosum.

The present study did not find any significant sex differences in the baboon corpus callosum: neither did we detect a main effect of sex on midsagittal area or thickness measures, nor did we find any indication for differences in the lifespan trajectories between the sexes. In general, sex difference in the corpus callosum have been frequently examined in various primate species and yield inconsistent results (e.g., rhesus monkeys: Franklin et al., 2000; Payne, Cirilli, & Bachevalier, 2017; marmosets: Sakai et al., 2017; capuchin monkeys: Phillips, Sherwood, & Lilak, 2007; chimpanzees: Hopkins et al., 2016; Phillips & Hopkins, 2012; Westerhausen et al., 2020). Best studied regarding sex differences is obviously the human corpus callosum. Here, meta-analytic evidence suggest relative corpus callosum area to be larger in females compared to males (Smith, 2005). The effect size is, however, rather small yielding a Cohen’s *d* of 0.2 (i.e., ca. 1% explained variance). Likewise studies examining callosal thickness in humans, while revealing significantly thicker female corpora callosa in some subregions (Danielsen et al., 2020; Luders, Toga, & Thompson, 2014; Smith, 2005), the effect size is usually small explaining below one percent of the variance. The sensitivity power analysis of the present study (see Supplement Section 6) indicates sufficient test power (>.80) to detect effect sizes of ca. 7% explained variance. Sex effects larger than this can thus be reliably excluded both concerning area and thickness measures. However, taken the human findings as “best guess” for what to expect in baboons, the present study would be underpowered. It remains for future studies to determine whether small sex differences exist in the baboon corpus callosum.

Finally, the interpretation of the present findings has to consider a set of limitation. Firstly, the present data analysis utilized cross□ sectional data so that the found aging trajectories reflect inter-individual differences between animals of different age rather than intra-individual changes. Consequently, potential differences between birth cohorts cannot be distinguished from age□ related changes on an individual level (Pfefferbaum & Sullivan, 2015; Salthouse, 2011). Thus, a confirmation in future mixed-effect or longitudinal studies is warranted. Secondly, the use of macroanatomical measures of the corpus callosum will not fully capture known age-related alteration on microstructural level. Histological and high-gradient strength diffusion-imaging studies demonstrate a decline in number and density of small myelinated axons (Fan et al., 2019; Hou & Pakkenberg, 2012; Køster et al., 2018) as well as a degeneration of axonal myelin sheaths (Bowley et al., 2010; Peters & Sethares, 2002) in old aged human and other primates. Combined histological and morphological analyses also report that the midsagittal area is a good predictor of the number of myelinated axons in the corpus callosum (Hou & Pakkenberg, 2012; Riise & Pakkenberg, 2011). Thus, we believe that any substantial deterioration of callosal axons in old age should also have been detected by the present area or thickness measures. Nevertheless, further histological or diffusion-imaging studies are required to examine callosal aging on an axonal level, and examine the reasons for differential aging trajectories between baboons, humans, and other primate species.

In summary, the baboon corpus callosum area follows a linear trajectory across the lifespan. Compared with other primate species, the trajectory is characterised by a slower increase from childhood into adulthood. Like all other non-human primates studied to-date, baboons do not show any signs of callosal atrophy in old age. In this, the present findings lend support to the hypothesis that atrophy of the corpus callosum is a unique characteristic of human brain aging. It remains for to explore the alterations in axon or myelin architecture, which underlie these divergent aging trajectories of corpus callosum morphology of humans and non-human primates.

## Supporting information

Supplemetal Figures and Analyses

## Acknowledgments

A.M has received funding from the European Research Council (ERC) under the European Union’s Horizon 2020 research and innovation programm grant agreement No 716931 (716931 - GESTIMAGE - ERC-2016-STG), from the French “Agence Nationale de le Recherche” (ANR-12-PDOC-0014-01, LangPrimate Project) as well as from grants ANR-16-CONV-0002 (ILCB) and the Excellence Initiative of Aix-Marseille University (A*MIDEX).

We are very grateful to the MRI Center of the INT in Marseille, Muriel Roth, Bruno Nazarian and Jean-Luc Anton. We thank also the Station de Primatologie CNRS of Rousset, particularly the vets Romain Lacoste, Ivan Balansard and Alice Bertello, the animal care staff and technicians: Jean-Noël Benoit, Jean-Christophe Marin, Valérie Moulin, Fidji and Richard Francioly, Laurence Boes, Célia Sarradin, Brigitte Rimbaud, Sebastien Guiol, Georges Di Grandi and Pau Molina.

R.W. conceptualized the study, analysed the data, and darfted the manuscript; A.M. provided the data and contributed to the discussion and interpretation of the findings

