## Supplementary material for "Corpus callosum morphology across the lifespan in baboons (*Papio anubis*): a cross-sectional study of relative mid-sagittal surface area and thickness": Supplemetal Figures and Analyses

**1. Figure S1: Mean outline of the baboon corpus callosum, geometric subdivision, and thickness segments**

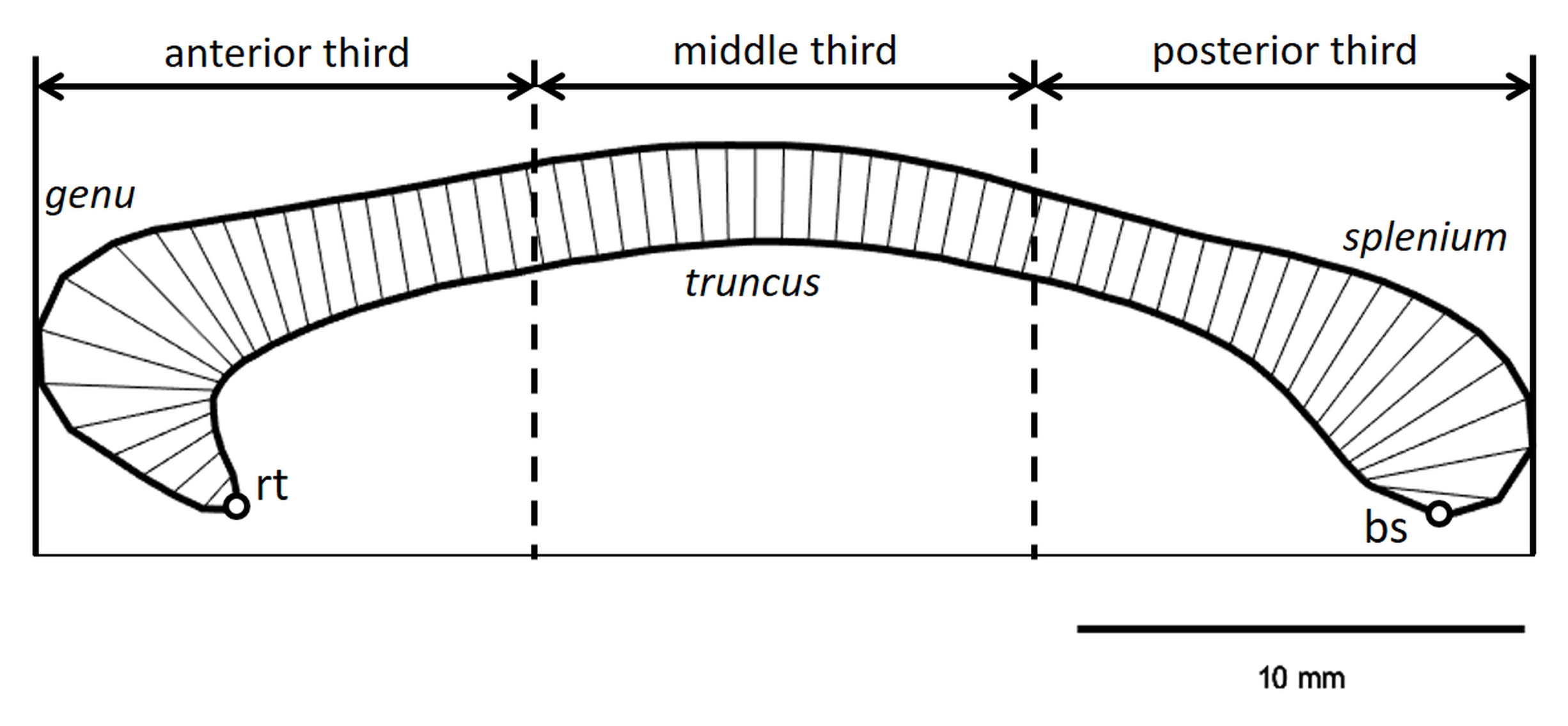

*Figure S1*. Mean outline of the corpus callosum across all subjects (scale: lower right corner). The points marked “rt” and “bs” denote the rostral tip and the bottom of the splenium, respectively. The imagined line between these two reference points was rotated into horizontal orientation for each individual to obtain a comparable reference across subjects. The figure additionally illustrates the geometric subdivision into thirds relative to anterior-posterior extend of the corpus callosum, which was utilized for the subsection analysis. The lines within the corpus callosum represent the 60 thickness measurement segments.

**2. Example R code for main GAM analyses**

Aging trajectories of relative area and thickness measures were fitted using the “mgcv” package (v1.8‐31; (Wood, 2017)) in R 3.6.2, using the following code template:

> gam.object <- gam(DV ~ s(Age, bs = "cr", k=4) + Sex, data = dataset, method = "REML")

Whereby DV was the respective dependent measures (e.g., relative area, relative subregional area or relative segmental thickness).

Sex differences in the aging trajectories were tested in a follow-up analysis incorporating terms for the modulation of the aging trajectories by sex. That is, the female age trajectory was fitted as reference, and deviation of the male trajectory from this reference trajectory was tested. Expressed as R code:

> gam.object <- gam(DV ~ s(Age, bs = "cr", k=4) + s(Age, bs = "cr", by= as.ordered(Sex)) + Sex, data = dataset, method = "REML")

**3. Analysis of absolute corpus callosum measures and Fig. S2**

The findings of absolute area measures mirrored the findings observed when analysing the relative measures. That is, analysing the total absolute corpus callosum area a significant effect of Age (see Fig S2 below; *F* = 3.70, edf = 1.17, *p* = .035, ω² = .04) was detected. Analysed by subregion, the Age effect was significant only in the anterior third (edf = 1.16, *F* = 4.13, *p* = .03, ω² = .04). For middle (edf = 1.00, *F* = 3.87, *p* = .052, ω² = .03) and posterior third (edf = 1.67, *F* = 2.60, *p* = .08, ω² = .03) the age effect might be seen as trend.

The main effect of Sex was neither significant for the total corpus callosum (*t*(87.8) = 0.95, *p* = 0.35, ω² < .01) nor any of the subregions (anterior: *t*(87.8) = 1.60, *p* = 0.11, ω² = .02; middle: *t*(88.0) = -0.12, *p* = 0.90, ω² < .01 , posterior: *t*(87.2) = 0.94, *p* = 0.35, ω² < .01). Neither for total area nor subregions sex differences in age trajectories were found (all *F* <1.86, all *p* > .14, all ω² < .04).

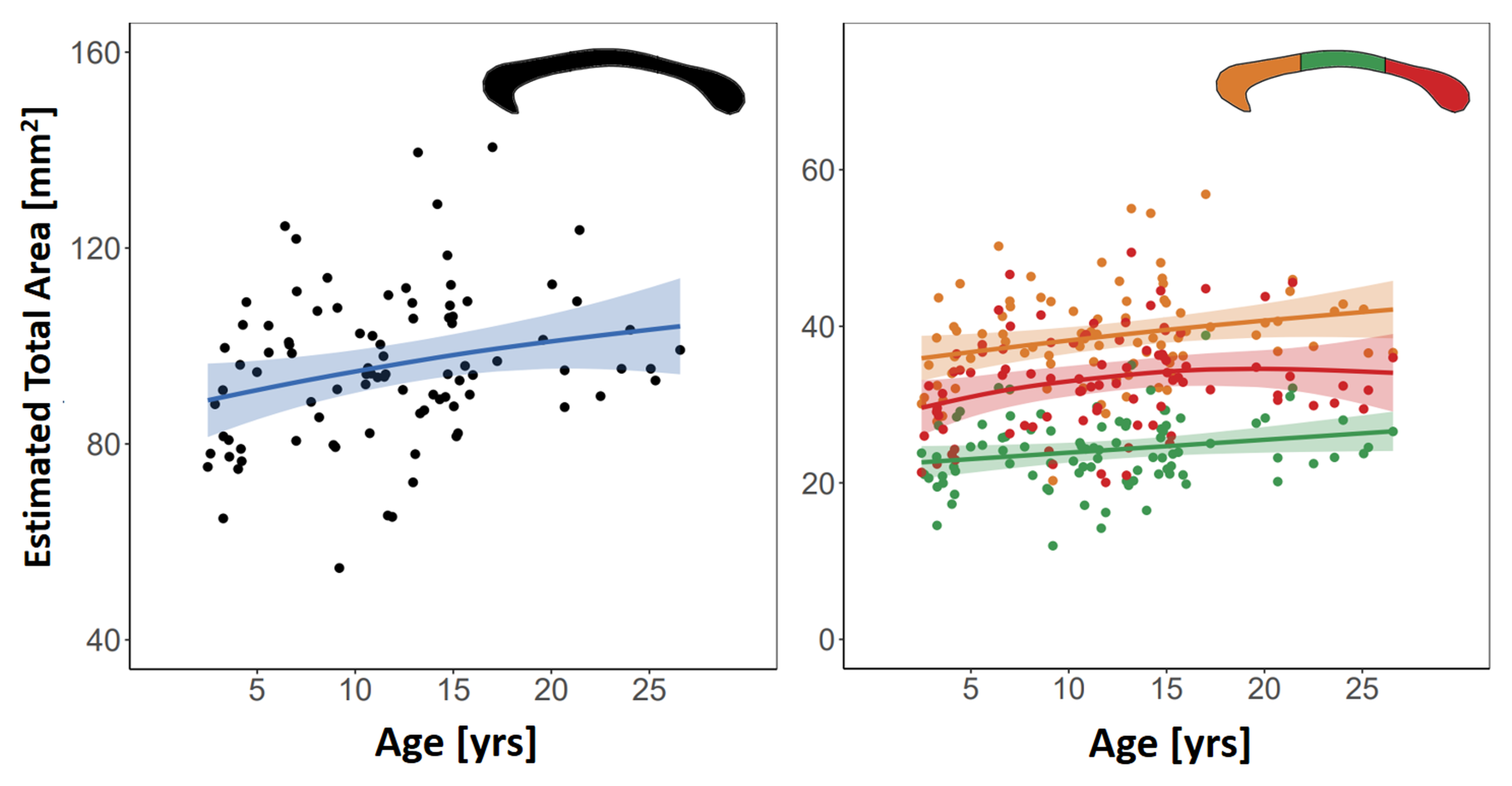

*Figure S2*. Association of age and corpus callosum area (left panel: total area, right panel: subsection area). The graphs show the GAM fitted age trajectories (shaded area 95% confidence bands).

The segment-wise analysis of absolute thickness, did not find any significant effect of Age (all edf < 2.00, all *p*_FDR_ >.72; all *ω*² < .11) or Sex (all *p*_FDR_ >.38; all *ω*² < .10).

**4. Table S1. Segment-wise thickness analysis**

| *Table S1* Results of the main and follow-up GAM analyses of relative thickness per segment | | | | | | | | |
| --- | --- | --- | --- | --- | --- | --- | --- | --- |
|  | Age trajectory fit | | | | Sex effect | | Deviation trajectory between Sexes | |
| # | *F* | edf | *p* | ev | *p* | ev | *p* | ev |
| 1 | 0.00 | 1.00 | 0.99 | 0.00 | 0.24 | 0.00 | 0.46 | 0.00 |
| 2 | 0.00 | 1.00 | 0.98 | 0.00 | 0.13 | 0.01 | 0.29 | 0.02 |
| 3 | 0.55 | 1.00 | 0.46 | 0.00 | 0.31 | 0.00 | 0.37 | 0.01 |
| 4 | 0.48 | 1.00 | 0.49 | 0.00 | 0.39 | 0.00 | 0.75 | 0.00 |
| 5 | 0.87 | 1.00 | 0.35 | 0.00 | 0.59 | 0.00 | 0.49 | 0.00 |
| 6 | 0.61 | 1.36 | 0.39 | 0.00 | 0.74 | 0.00 | 0.28 | 0.00 |
| 7 | 1.05 | 1.03 | 0.30 | 0.00 | 0.53 | 0.00 | 0.09 | 0.05 |
| 8 | 1.86 | 1.00 | 0.18 | 0.01 | 0.14 | 0.01 | 0.06 | 0.07 |
| 9 | 1.65 | 1.00 | 0.20 | 0.01 | 0.10 | 0.02 | 0.79 | 0.00 |
| 10 | 3.58 | 1.00 | 0.06 | 0.03 | 0.51 | 0.00 | 0.54 | 0.00 |
| 11 | 0.08 | 1.00 | 0.77 | 0.00 | 0.18 | 0.01 | 0.68 | 0.00 |
| 12 | 1.89 | 1.00 | 0.17 | 0.01 | 0.96 | 0.00 | 0.28 | 0.00 |
| 13 | 3.61 | 1.00 | 0.06 | 0.03 | 0.80 | 0.00 | 0.07 | 0.03 |
| 14 | 3.16 | 1.00 | 0.08 | 0.02 | 0.97 | 0.00 | 0.28 | 0.00 |
| 15 | 3.32 | 1.00 | 0.07 | 0.02 | 0.45 | 0.00 | 0.90 | 0.00 |
| 16 | 2.62 | 1.31 | 0.15 | 0.03 | 0.12 | 0.02 | 0.68 | 0.00 |
| 17 | 2.30 | 1.00 | 0.13 | 0.01 | 0.07 | 0.03 | 0.56 | 0.00 |
| 18 | 3.92 | 1.00 | 0.05 | 0.03 | 0.46 | 0.00 | 0.33 | 0.00 |
| 19 | 3.82 | 1.00 | 0.05 | 0.03 | 0.65 | 0.00 | 0.41 | 0.00 |
| 20 | 3.61 | 1.00 | 0.06 | 0.03 | 0.18 | 0.01 | 0.86 | 0.00 |
| 21 | 4.02 | 1.00 | 0.048 | 0.03 | 0.15 | 0.01 | 0.99 | 0.00 |
| 22 | 1.29 | 1.00 | 0.26 | 0.00 | 0.01 | 0.07 | 0.00 | 0.19 |
| 23 | 3.30 | 1.00 | 0.07 | 0.02 | 0.05 | 0.03 | 0.59 | 0.00 |
| 24 | 2.78 | 1.00 | 0.10 | 0.02 | 0.04 | 0.04 | 0.76 | 0.00 |
| 25 | 2.03 | 1.00 | 0.16 | 0.01 | 0.00 | 0.08 | 0.69 | 0.00 |
| 26 | 2.16 | 1.00 | 0.14 | 0.01 | 0.01 | 0.06 | 0.12 | 0.04 |
| 27 | 0.93 | 1.00 | 0.34 | 0.00 | 0.12 | 0.02 | 0.07 | 0.06 |
| 28 | 0.39 | 1.00 | 0.53 | 0.00 | 0.20 | 0.01 | 0.44 | 0.00 |
| 29 | 0.05 | 1.00 | 0.83 | 0.00 | 0.05 | 0.03 | 0.27 | 0.02 |
| 30 | 0.09 | 1.20 | 0.75 | 0.00 | 0.10 | 0.02 | 0.75 | 0.00 |
| 31 | 0.28 | 1.00 | 0.60 | 0.00 | 0.27 | 0.00 | 0.81 | 0.00 |
| 32 | 0.00 | 1.00 | 1.00 | 0.00 | 0.16 | 0.01 | 0.17 | 0.03 |
| 33 | 2.61 | 1.00 | 0.11 | 0.02 | 0.01 | 0.07 | 0.00 | 0.21 |
| 34 | 0.09 | 1.00 | 0.77 | 0.00 | 0.09 | 0.02 | 0.26 | 0.00 |
| 35 | 0.09 | 1.00 | 0.76 | 0.00 | 0.08 | 0.02 | 0.28 | 0.01 |
| 36 | 0.02 | 1.00 | 0.88 | 0.00 | 0.02 | 0.05 | 0.36 | 0.00 |
| 37 | 0.93 | 1.00 | 0.34 | 0.00 | 0.08 | 0.02 | 0.78 | 0.00 |
| 38 | 1.05 | 1.00 | 0.31 | 0.00 | 0.15 | 0.01 | 0.96 | 0.00 |
| 39 | 0.91 | 1.00 | 0.34 | 0.00 | 0.27 | 0.00 | 0.99 | 0.00 |
| 40 | 1.45 | 1.00 | 0.23 | 0.00 | 0.12 | 0.02 | 0.69 | 0.00 |
| 41 | 1.02 | 1.31 | 0.25 | 0.00 | 0.42 | 0.00 | 0.46 | 0.00 |
| 42 | 0.72 | 1.23 | 0.34 | 0.00 | 0.78 | 0.00 | 0.47 | 0.00 |
| 43 | 0.80 | 1.57 | 0.36 | 0.00 | 0.48 | 0.00 | 0.16 | 0.03 |
| 44 | 2.86 | 1.58 | 0.09 | 0.04 | 0.18 | 0.01 | 0.97 | 0.00 |
| 45 | 0.65 | 1.30 | 0.36 | 0.00 | 0.43 | 0.00 | 0.36 | 0.01 |
| 46 | 0.93 | 1.53 | 0.30 | 0.00 | 0.83 | 0.00 | 0.21 | 0.03 |
| 47 | 2.41 | 1.87 | 0.08 | 0.03 | 0.31 | 0.00 | 0.11 | 0.05 |
| 48 | 1.96 | 1.98 | 0.11 | 0.02 | 0.87 | 0.00 | 0.34 | 0.01 |
| 49 | 2.25 | 1.86 | 0.10 | 0.03 | 0.72 | 0.00 | 0.24 | 0.02 |
| 50 | 2.39 | 1.72 | 0.09 | 0.03 | 0.56 | 0.00 | 0.24 | 0.02 |
| 51 | 4.07 | 1.81 | 0.02 | 0.07 | 0.49 | 0.00 | 0.67 | 0.00 |
| 52 | 3.87 | 1.83 | 0.02 | 0.06 | 0.89 | 0.00 | 0.92 | 0.00 |
| 53 | 5.80 | 1.92 | 0.00 | 0.11 | 0.97 | 0.00 | 0.76 | 0.00 |
| 54 | 4.98 | 1.83 | 0.01 | 0.09 | 0.89 | 0.00 | 0.76 | 0.00 |
| 55 | 2.16 | 1.00 | 0.14 | 0.01 | 0.60 | 0.00 | 0.87 | 0.00 |
| 56 | 4.18 | 1.68 | 0.02 | 0.07 | 0.73 | 0.00 | 0.58 | 0.00 |
| 57 | 1.53 | 1.00 | 0.22 | 0.01 | 0.34 | 0.00 | 0.92 | 0.00 |
| 58 | 2.67 | 1.00 | 0.11 | 0.02 | 0.52 | 0.00 | 0.66 | 0.00 |
| 59 | 0.70 | 1.79 | 0.44 | 0.00 | 0.60 | 0.00 | 0.10 | 0.07 |
| 60 | 0.14 | 1.00 | 0.71 | 0.00 | 0.45 | 0.00 | 0.28 | 0.02 |
| *Notes*. All p-values are uncorrected, and no effect survived FDR correction (to 5%). “ev” explained variance | | | | | | | | |

**5. Additional information to Figure 3**

Both panels of Figure 3 (main manuscript) provide a visual comparison of lifespan trajectories of relative total corpus callosum area of baboons, humans, and chimpanzees. While the baboon data stems from the present manuscript, the human and chimpanzee data was taken from (Westerhausen et al., 2020). The graphs shows the GAM aging trajectories fitted separately for the three species using the “mgcv” R-package (v1.8‐31; (Wood, 2017); R build: 3.6.2). Cubic regression splines with k=6 knots as basis dimension was used for all three species to use the same parameters for all fits. The left panel shows the comparison after correcting for differences in longevity between the three species. That is, the age of baboons and chimpanzees was transferred to a human age equivalent using multiplication factors suggested in the literature. More specifically, the age of baboons was multiplied with a factor of 3.25 (as factors between 3 and 3.5 have been suggested; see e.g. (Franke et al., 2017; Havill et al., 2005)), the age of chimpanzees with a factor of 1.5 (for discussion see Westerhausen et al., 2020). The right panel shows the same comparison for chronological age. For both panel, the data of each species was divided by the relative corpus callosum area predicted for the age of 2 years (human age equivalent) using the GAM fitted for the respective species. Thus, the value 1 represent the predicted relative area at the age of 2 years. The age of 2 years rather than 0 years (birth) was chosen as reference age as the corpus callosum is known to grow rapidly during infancy and the growth rates differ substantially between the species (Phillips & Kochunov, 2011; Sakai et al., 2017; Vannucci, Barron, & Vannucci, 2017). This differences would, however, not be reflected in the GAM prediction as none of the samples include individuals below the age of 2 years.

**6. Sensitivity power analysis for the main effect of sex**

The power analysis for the main effect of Sex was conducted using GPower (version 3.1.9.2; (Faul, Erdfelder, Buchner, & Lang, 2009)). The “generic t-test” function to be able to account for the fact that the main effect of Sex was embedded in the GAM model and the degrees of freedom of the t-test were accordingly adjusted. GPower provides the following summary:

**t tests -** • Generic t test

**Analysis:** Sensitivity: Compute noncentrality parameter

**Input:** Tail(s)	=	One

α err prob	=	0.05

Power (1-β err prob)	=	0.80

Df	=	88

**Output:** Critical t	=	1.6623540

Noncentrality parameter δ	=	2.5058151

To convert the obtained noncentrality parameter *δ* to Cohen’s *d* under consideration of that the sample sizes *n_f_* = 57 and *n_m_* = 34 for females and males, respectively, we used the following formula (source: GPower manual):

$$d=\frac{\delta}{\sqrt{\frac{n_{f}n_{m}}{n_{f}+n_{m}}}}= \frac{2.5058}{4.6148}=0.5430$$

The resulting *d* can be transformed to proportion of explained variance (*ev*) as follows:

$$ev=\frac{d^{2}}{d^{2}+4}=\frac{0.2948}{4.2948}=0.0687$$
